# Transcriptomic Profiling of scWAT Reveals Reduced Lipid-Associated Macrophage Signatures in Long-Term Endurance- and Resistance-Trained Athletes

**DOI:** 10.1101/2025.11.07.687158

**Authors:** Eric B. Emanuelsson, Stefan M. Reitzner, Allister Quizon, Joshua T. Burrows, Merve Elmastas, Jutta Jalkanen, Sean Perez, Hampus Gilljam, Malene E Lindholm, Jessica Norrbom, Bernhard O. Palsson, Daniel C. Zielinski, Mikael Rydén, Mark A. Chapman, Carl Johan Sundberg

**Author notes:** These authors have contributed equally (Eric B. Emanuelsson, Stefan M. Reitzner). These authors jointly supervised this work (Mark A. Chapman, Carl Johan Sundberg).

## Abstract

Regular exercise training is associated with systemic and tissue-specific performance and health benefits, yet long-term molecular adaptations in subcutaneous abdominal white adipose tissue (scWAT) remain unclear. We analyzed the resting scWAT transcriptome of 89 adults (35–50 years) who were long-term (>15 years) endurance- or strength-trained, or untrained. Long-term training was linked to enrichment of ribosomal, mitochondrial, and aerobic metabolism-related pathways, while immune-related pathways were enriched in untrained individuals. A distinct sex dimorphic scWAT was identified: untrained females exhibited a more metabolically favorable, ‘trained-like’ transcriptomic profile, and untrained males an ‘untrained-like’ profile. In trained males, the inflammatory gene signature was lower than in untrained males. We also identified 11 genes specific to lipid-associated macrophages with lower expression in trained individuals, particularly in all endurance-trained individuals and strength-trained females. These findings highlight sex-specific, exercise-induced adaptations in scWAT relevant to metabolic health, and underscore the importance of including both sexes in scWAT research.

## Introduction

White adipose tissue (WAT) plays a pivotal role in energy homeostasis, acting as a primary site for lipid storage and an active endocrine organ^1^. The dynamic nature of WAT allows it to respond to various physiological stimuli, including nutritional status, hormonal signals, and physical activity^2,3^. Exercise training, a well-established modulator of metabolic health, exerts profound effects on WAT by reducing WAT mass, enhancing lipolysis and improving insulin sensitivity of glucose uptake in subcutaneous WAT (scWAT)^4–6^. However, the molecular mechanisms behind WAT adaptations to regular training are incompletely understood. Most previous studies have investigated short-term training interventions^3,7–10^, while studies of individuals with several years of regular training are limited^11,12^. To fully understand the health and performance benefits acquired over time, studies with long-term trained individuals are needed. Moreover, while most studies on scWAT to date have focused on endurance training^7–10^ or combined endurance and strength protocols^5,13^, investigations of isolated strength training are scarce^14,15^, and studies of long-term strength-trained males or females are lacking.

Previous studies examining the effects of exercise on scWAT have either excluded one sex^5,8–10,16^ or failed to analyze males and females separately^11,17,18^. In studies addressing baseline differences in untrained individuals, it has been shown that scWAT is one of the most sexually dimorphic tissues in humans^19,20^, with males generally having a larger proportion of hypertrophic adipocytes, which has been linked to increased inflammation^21^. A recent study in rats found a sexually dimorphic scWAT transcriptomic signature in sedentary animals which was maintained or amplified following endurance training^22^. Biological processes related to insulin signaling and adipogenesis were enriched in female rats while male rats were enriched for aerobic metabolism^22^. Although mechanistic studies in animal models provide valuable insights to improve our understanding of adaptations to various stressors, a previous integrative analysis of gene expression and gene set enrichment showed that few differentially expressed genes (DEGs) and Gene Ontology (GO) terms overlapped between rodent and human white adipose tissue^5^. Thus, there is a need for sex-specific comparisons of scWAT global gene expression in long-term trained and untrained humans.

Here, we recruited 89 individuals (47 females, 42 males) into three experimental groups: long-term (>15 years) endurance-trained athletes, long-term strength-trained athletes, and age-matched untrained individuals with a body mass index (BMI) ≤ 27. This design enabled us to examine the impact of long-term training on the scWAT transcriptome as well as sex differences between these groups. Finally, using public datasets, we compared training-associated DEGs to those observed in individuals with type 2 diabetes.

## RESULTS

### Large differences in adipose tissue gene expression between endurance trained and untrained individuals

A flowchart of the study design is presented in Figure 1A. In short, we recruited 89 individuals (47 females, 42 males) into three experimental groups: long-term (>15 years) endurance-trained athletes (20 females [FE] and 13 males [ME]), long-term strength-trained athletes (9 females [FS] and 14 males [MS]), and untrained healthy age-matched individuals as controls (18 females [FC] and 15 males [MC]). Subject characteristics are presented in Figure 1B & 1C. For further details on the recruitment process, see methods. By design, the endurance groups had significantly higher VO_2_-peak than the other groups, while the strength groups had significantly higher peak torque than the other groups (Figure 1C).

**Figure 1.**
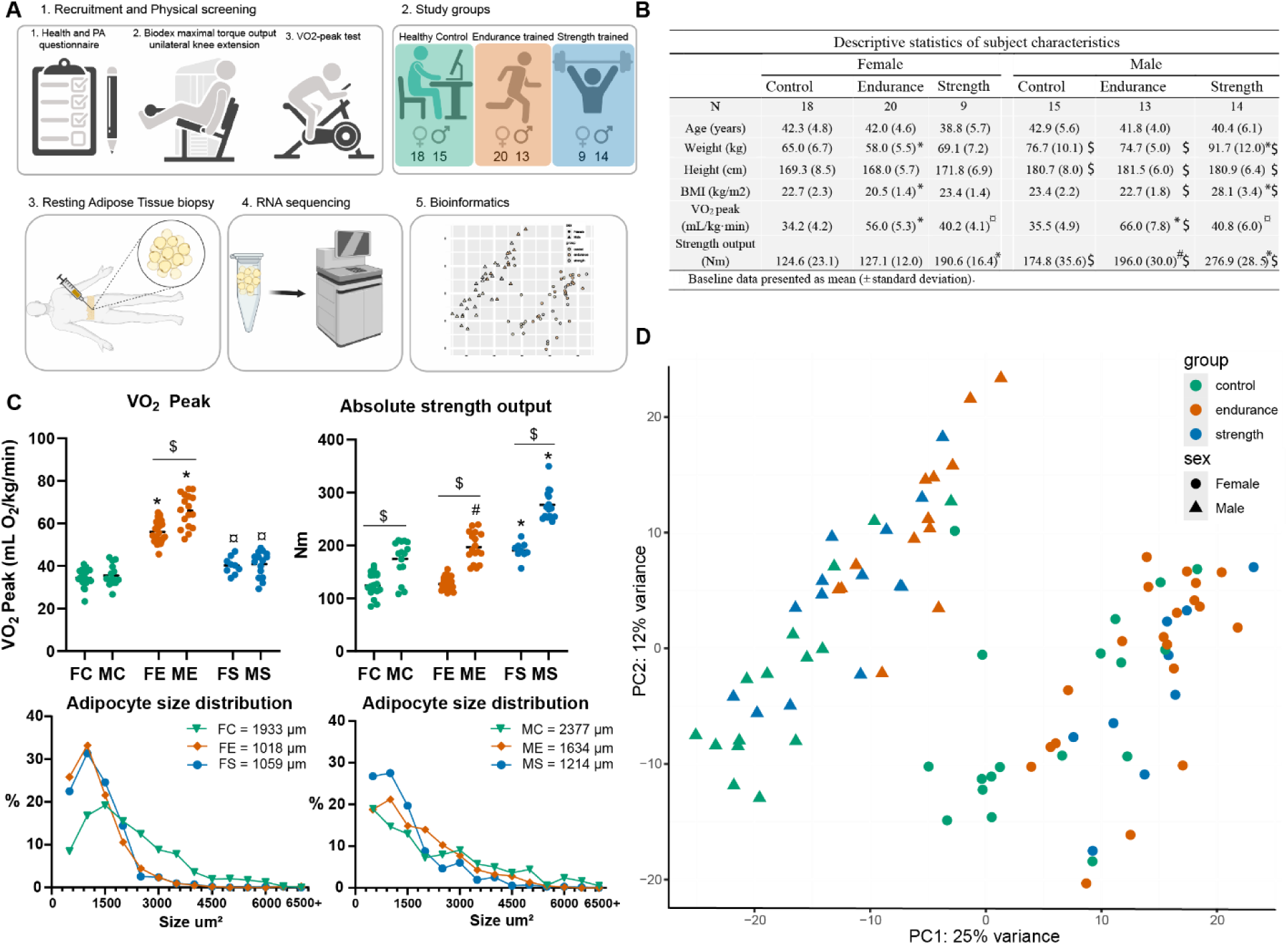
Study design and subject characteristics A) Flowchart of the study design. B) Subject characteristics. C) VO_2peak_, absolute strength output and adipocyte size distribution in the three groups. The average adipocyte size value is indicated beside each group abbreviation. D) Principal component analysis displayed by exercise background and sex. * = significantly different to the other sex-matched groups, $ = significant sex-difference within an experimental group. # = significantly different between sex-matched endurance and control groups, ¤ = significantly different between sex-matched strength and control groups. F = female, M = male, C = control, E = endurance, S = strength. Created in BioRender (Icons in Figure 1A). Emanuelsson, E. (2025) https://BioRender.com/7za25n8.

Abdominal scWAT biopsies were collected and adipocyte size was determined histologically on a subset of the samples (three samples from each training background and sex). Trained subjects showed 31-49% smaller average adipocyte areas and a higher proportion of smaller adipocytes compared with the untrained groups (Figure 1C). Following the confirmation of adipocyte sizes, RNA sequencing was performed on all samples. A principal component analysis (PCA) of the scWAT transcriptome showed separation based on sex (PC1) and training background (PC2), with the separation between sexes being more prominent (Figure 1D). Additionally, the separation observed based on training background was more pronounced in the male cohort than in the female cohort (Figure 1D).

#### Endurance trained and untrained individuals display distinctly different scWAT gene expression

Differential expression analysis identified the largest training-induced differences to be between endurance-trained and untrained in both female (2070 DEGs: 821 with higher and 1249 with lower expression in FE) and male participants (2417 DEGs: 1150 with higher and 1267 with lower expression in ME; Figure 2A, Supplemental Data 1). Of these, 246 of the higher expressed and 467 of the lower expressed DEGs were shared by both ME and FE compared with MC and FC, respectively. In these shared DEGs, only one gene, *POLR2H*, was oppositely regulated between ME (up) and FE (down). Among the genes with the largest fold-difference (positive or negative) between FE vs FC and ME vs MC, the endurance trained groups in both sexes displayed lower expression of the *EGFL6* and *TM4SF19* genes and a higher expression of *NTSR2* (Figure 2A).

**Figure 2:**
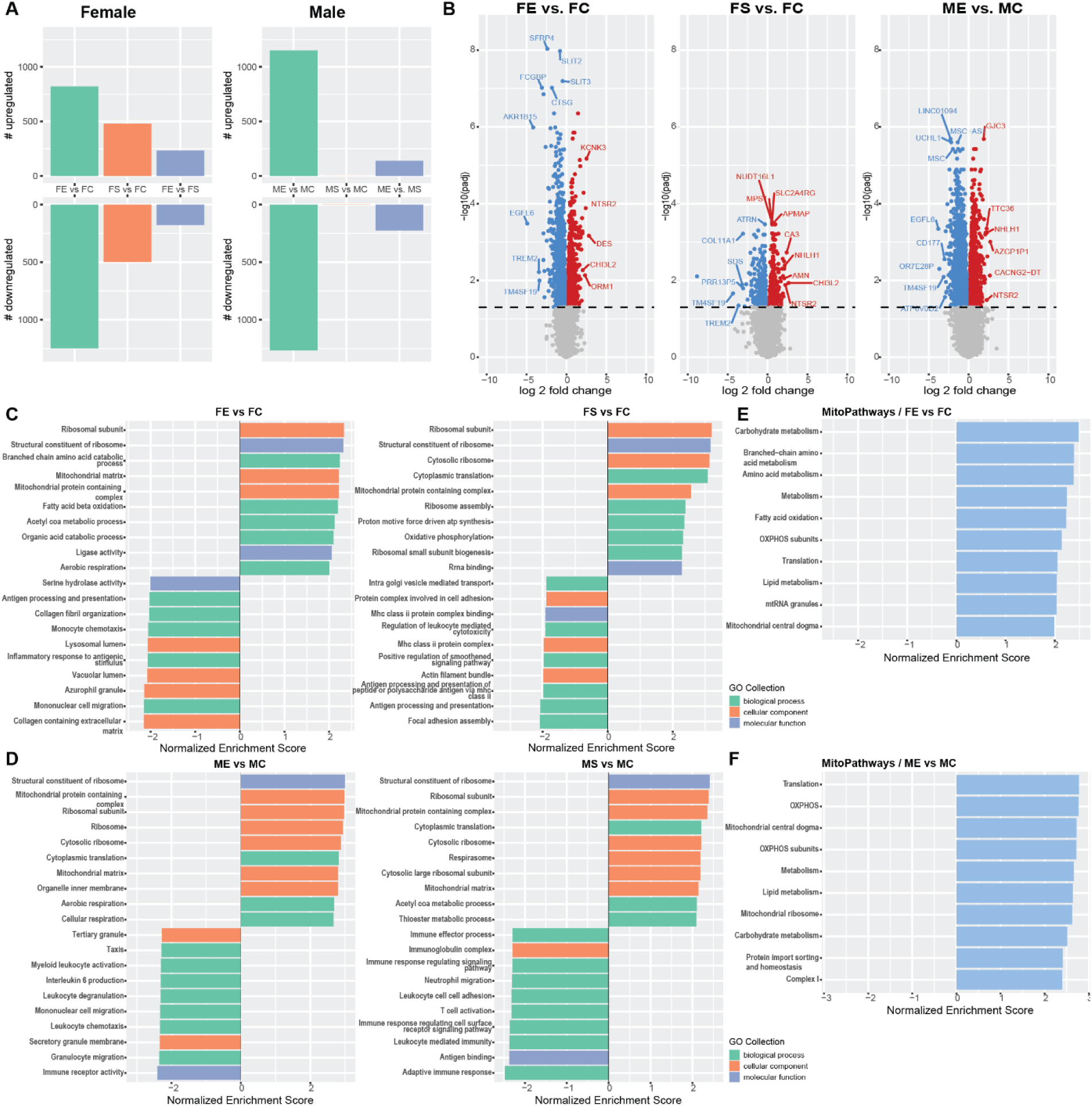
Differences in gene expression between trained and untrained individuals. A) Number of differentially expressed genes between trained and untrained females and males, respectively. B) Volcano plots with annotations of the top five most significantly expressed genes, the five genes with highest and lowest fold differences between groups. C-D) Fast gene set enrichment analysis (fGSEA) with GO-terms displaying the top 10 terms with the highest or lowest normalized enrichment score in each comparison. E-F) fGSEA of MitoPathways. F = female, M = male, C = control, E = endurance, S = strength, GO = Gene Ontology, padj = adjusted p-value.

To better understand the functional differences between the endurance trained and control groups, we performed Gene Set Enrichment Analysis (GSEA) using the GO collections. GSEA results showed that the top 10 most enriched terms in both ME and FE compared to their respective control group were related to ribosomal, aerobic energy metabolism and mitochondrial terms (Figure 2C-D). Comparing ME vs MC, 8 of the top 10 most enriched GO terms in MC were related to the immune system. This enrichment pattern was also visible comparing FE vs FC, with the addition of extracellular matrix and lysosomal terms being present in the top 10 most enriched terms in FC (Figure 2C-D).

Considering the large number of significantly enriched mitochondrial terms in the endurance-trained groups, we further investigated mitochondria-specific annotations using the MitoPathway annotations^23^. All pathways significantly enriched in this analysis were higher in ME and FE compared with MC and FC, respectively (Figure 2E-F). Interestingly, substantially more pathways were enriched in ME compared with MC than in their female counterparts (ME vs MC n = 76 compared with FE vs FC n = 25; Supplemental Data 1). Comparing the top 10 terms in ME vs MC and FE vs FC showed that 6/10 were common to both comparisons which included the mitochondrial central dogma, OXPHOS subunits and carbohydrate metabolism pathways (Figure 2E-F).

#### Strength trained females, but not males, display a large number of DEGs compared with control group

Comparing the female and male strength groups with their respective control group, we found 976 DEGs (479 with higher and 497 with lower expression) between FS and FC while only two DEGs were identified in the male counterparts (Figure 2A). However, GSEA showed similar patterns in both comparisons with GO terms related to the ribosome, mitochondria and aerobic energy metabolism among the top 10 most highly enriched terms (Figure 2C-D). In the control groups, both MC and FC displayed immune system-related terms among the top 10 most enriched (up in MC and FC, respectively, Figure 2C-D). Similarly to the endurance groups, both FS and MS had mitochondrial carbohydrate metabolism OXPHOS subunits and translation pathways among the top 10 most enriched mitochondria-specific pathways when compared to FC and MC (Supplemental Data 1).

#### Training background-specific transcriptome in endurance- and strength-trained athletes

As long-term endurance and strength training induce distinct whole-body and tissue-specific adaptations^24–27^, we aimed to investigate whether training background influences scWAT transcriptomic profiles by directly comparing endurance-trained and strength-trained individuals. Comparing FE vs FS and ME vs MS, a total of 362 (139 with higher and 223 with lower expression) and 409 (233 with higher and 176 with lower expression) genes were differentially expressed, respectively (Figure 2A). In the male subjects, an enrichment of mitochondrial and ribosomal terms was found among the top 10 terms in ME, while MS displayed an enrichment of immune system related terms (Supplemental Data 1). When comparing FE vs FS, the top enriched terms in FE were helicase activity and ATP hydrolysis activity, while the top terms in FS were dominated by ribosomal GO-terms (Supplemental Data 1). Furthermore, when investigating mitochondrial pathways using MitoPathway, a substantially larger number of pathways were differentially enriched between the male groups (n = 75) than between the female groups (n = 1; Supplemental Data 1). In ME, mitochondrial translation, OXPHOS subunits and carbohydrate metabolism were the most enriched pathways compared with MS, while the enrichment of nucleotide import in FE was the only pathway significantly different compared with FS.

#### Lower inflammatory scWAT profile in trained compared to untrained males

Considering the substantial enrichment of inflammatory GO terms in the untrained males and females compared with both the endurance and strength trained groups, we used CIBERSORTx^28^ to infer the immune cell type proportions between groups based on exercise background. Compared with the male control group, both ME and MS displayed a lower inferred abundance of monocytes (Supplemental data 2). Furthermore, ME also displayed a lower abundance of M2 macrophages, neutrophils, and resting mast cells (Supplemental data 2). The male strength group had a lower abundance of M1 macrophages, CD8+ T-cells than MC. Both FE and FS displayed a lower M2 macrophage abundance than FC, and FE had lower resting mast cell expression than FC while FS had lower M0 macrophages than FC (Supplemental Data 2). For a full comparison of all cell populations, see Supplemental Data 2.

### Increased expression of genes related to aerobic energy metabolism in female scWAT

The differential gene expression analysis revealed more pronounced differences between sexes than between exercise groups. Specifically, 5244 genes were differentially expressed between ME and FE (2431 higher, 2813 lower), 3961 between MS and FS (1962 higher, 1999 lower), and 1810 between MC and FC (813 higher, 997 lower; Figure 3A & B).

**Figure 3.**
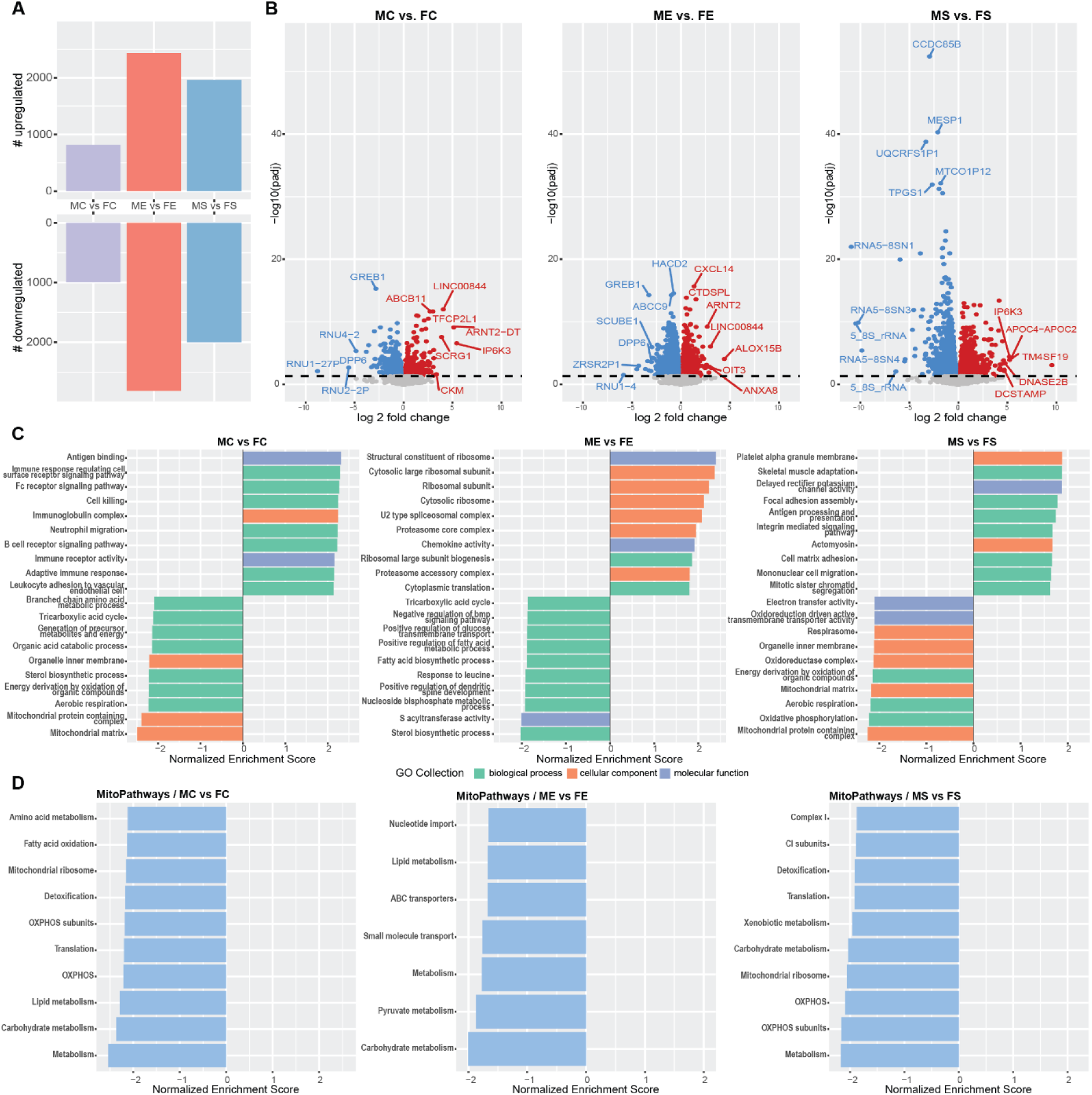
Sex differences in adipose tissue gene expression. A) Number of differentially expressed genes between males and females with similar exercise-background. B) Volcano plots with annotations of the top five most significantly expressed genes, the five genes with highest and lowest fold differences between groups. C) Fast gene set enrichment analysis (fGSEA) with GO-terms displaying the top 10 terms with the highest or lowest normalized enrichment score in each comparison. D) fGSEA of MitoPathways. F = female, M = male, C = control, E = endurance, S = strength, GO = Gene Ontology, padj = adjusted p-value.

Across all groups, 655 sex-specific DEGs (442 higher and 213 lower expressed in females) were identified. Overrepresentation analysis of genes with higher expression in females showed significant enrichment of the mitochondrial matrix and the inner and outer mitochondrial membrane. Genes with higher expression in males were enriched for immune-related processes, such as the leukocyte cell-cell adhesion and defense response to virus.

Next, to further explore sex differences in scWAT, GSEA was performed within each experimental group, revealing the highest number of significantly enriched GO terms between MC and FC (179 terms), followed by ME vs. FE (102 terms), and MS vs. FS (49 terms). In untrained males vs females, a distinct enrichment of immune system-related terms was found in males, whereas females showed enrichment in mitochondrial structure and aerobic respiration-related terms (Figure 3C). Similarly, trained females (FE and FS) exhibited enrichment in aerobic energy metabolism pathways, including the TCA cycle, fatty acid metabolism, oxidative phosphorylation, and electron transfer activity, compared to their male counterparts (Figure 3C). Following specific investigation of MitoPathways^23^, we found that females consistently show a positive enrichment in mitochondrial energy metabolism pathways, regardless of training status. The highest number of enriched mitochondrial pathways was observed in MC vs. FC (n = 45), followed by MS vs. FS (n = 27) and ME vs. FE (n = 7) (Figure 3D; Supplemental Data 1). All enriched pathways were positively enriched in females, and pathways related to mitochondrial carbohydrate and lipid metabolism were consistently enriched independent of exercise background. In contrast, trained males showed enrichment in ribosome-related (ME), while immune system-related and structural terms were enriched in MS (Figure 3C). Notably, no immune system-related GO terms were differentially enriched between ME and FE, unlike in the untrained group where numerous immune system-related terms showed differential enrichment (Supplemental Data 1).

Building on these findings, we examined the functions of genes exclusively differentially expressed in ME vs. MC or FE vs. FC to explore sex-specific training adaptations. Among the 800 DEGs with uniquely lower expressions in ME, we found significant enrichment of terms related to inflammatory response, positive regulation of cytokine production, and neutrophil migration. In contrast, the 781 DEGs with uniquely lower expressions in FE were enriched for biological processes such as cellular component disassembly, as well as extracellular matrix disassembly and organization. Meanwhile, the 903 genes with uniquely higher expression in ME were associated with enrichment of pathways related to the ribosome, mitochondrial translation, and cellular respiration. In ME however, the 575 genes with higher expression exclusive to FE revealed enrichment in cyclic-nucleotide-mediated signaling, cAMP-mediated signaling, and cholesterol metabolic processes.

#### Cell deconvolution analysis indicates an inflammatory profile of untrained male versus female scWAT

The untrained males displayed a significantly higher inferred abundance of monocytes, M1 macrophages, resting NK-cells, plasma cells, activated memory CD4+ T-cells and CD8+ T-cells compared with the untrained females (Supplemental Data 2), while no cell types were significantly higher in the FC group. Additionally, MS had higher expression of M2 macrophages than FS, and ME displayed higher expression of resting NK cells than FE, while both MS and ME had lower expression of resting memory CD4+ T-cells compared with FS and FE, respectively (Supplemental Data 2).

### Trained individuals have lower expression of genes enriched in lipid-associated macrophages

As pathway enrichment analyses are biased by pre-defined groups of genes that may not be suited to a tissue or species of interest, we additionally performed independent component analysis (ICA) to detect additional group-specific signatures within the data. ICA has been shown to effectively decompose and identify statistically independent components (ICs) of functionally relevant gene sets from human expression datasets^29–31^. Thus, ICA can be used to identify ICs to create unbiased tissue- and/or condition-specific signatures which can then be annotated to biological processes and functions.

The top five independent components together explained 45% of the variance, and two of these components showed a clear separation based on sex (ICs 17 and 18; Figure 4A, Supplemental data 3), while three separated by exercise background (ICs 3, 14 and 24; Figure 4A; Supplemental data 3). To confirm that biologically relevant components were detected by ICA, we first examined the sex-component IC 17, which included 11 Y-linked genes and the X-inactive specific transcript (XIST) gene (Supplemental Data 3). IC 18 also showed separation of the expression profiles across sex, but the genes involved in this component were not found on sex-linked chromosomes, indicating that they may be related to broader differences between sexes.

**Figure 4.**
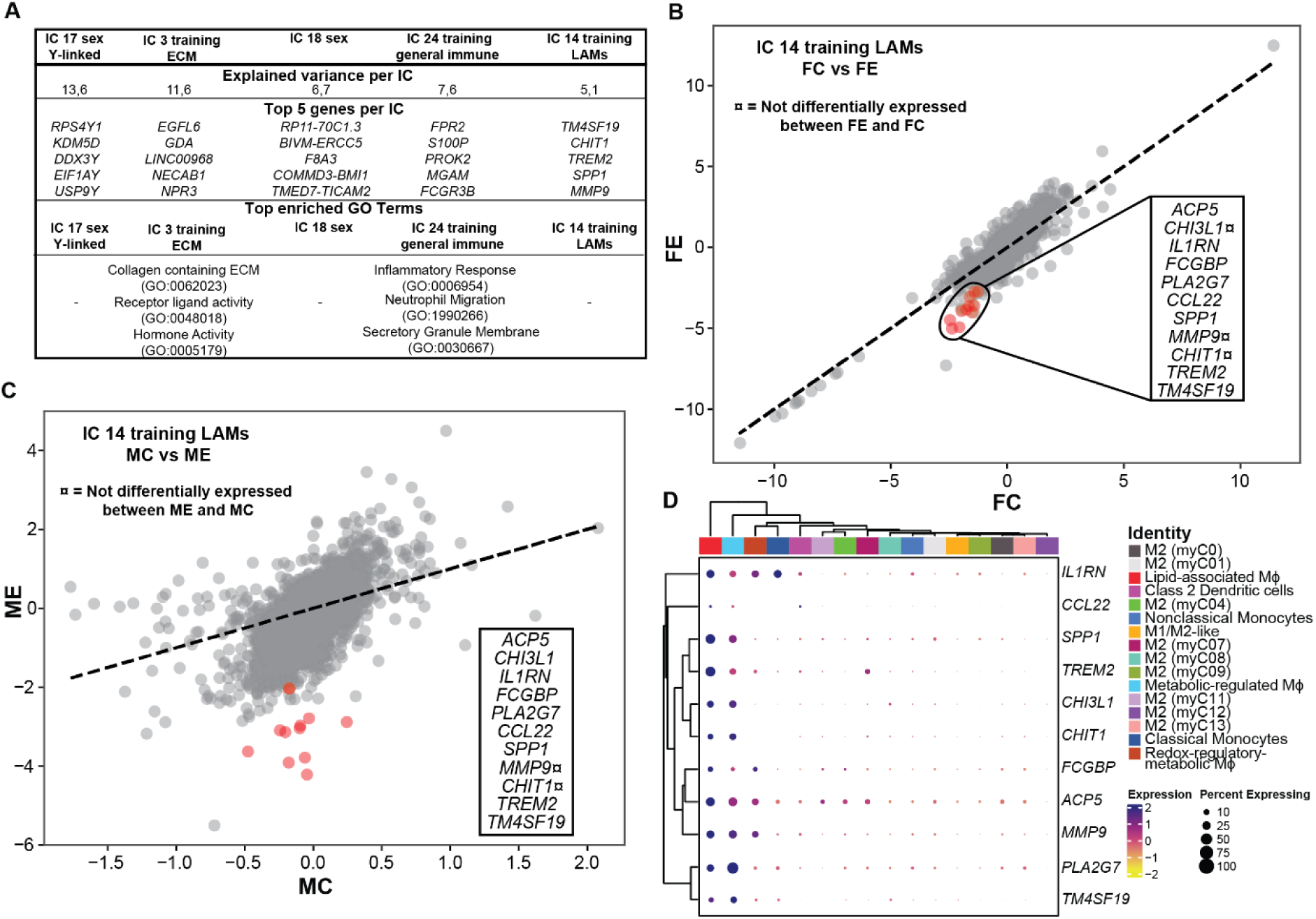
Independent component analysis. A) A descriptive table of the top five independent components (ICs) with the highest explained variance across all ICs, the top five genes with the highest weight in each IC and the top GO terms following overrepresentation analysis of the genes in the IC. Overrepresentation analysis of IC 17 and 14 was not performed due to the small number of genes in those ICs while the overrepresentation of IC 18 did not identify any statistically significant GO terms. B & C) Comparison of the average activity of all genes in IC14 between FC and FE, and MC vs MF respectively. The red circles highlight all genes in IC 14, and the gene names are displayed in the rectangle. ¤ Indicates that the gene was not identified as differentially expressed in the previous differential gene expression analysis, all other genes were differentially expressed. The dotted line is the 1:1 line and genes along that line have an identical activity in both groups. D) Clustered dot plot of all genes in IC 14 based on single-cell expression (using the dataset by Massier et al^33^) in various monocyte and macrophage (Mϕ) subpopulations. The M2-like (M2) subpopulations were named as in Massier et al^33^. Percent expressing indicates the proportion of cells within a cluster that express the specific gene. LAMs (Lipid-associated macrophages). F = female, M = male, C = control, E = endurance, S = strength, GO = Gene Ontology, padj = adjusted p-value.

Among the exercise-associated components, IC 3 showed the most distinct activity difference, indicating differential expression of genes within the component, between endurance-trained and untrained groups. Approximately 60% of IC 3 genes overlapped with differentially expressed genes from trained vs. control comparisons, indicating strong consistency with previous analyses. Overrepresentation analysis revealed enrichment in terms related to the extracellular matrix and receptor-ligand activity. Next, IC 24 was enriched for inflammatory response pathways, particularly neutrophil and granulocyte chemotaxis. It displayed lower activity in ME vs. MC, with 85% of its genes also being DEGs in the previous comparison, suggesting its relevance to training-induced inflammatory modulation. Moving on, IC 14 highlighted a specific set of 11 genes, of which 9 were DEGs in ME vs both MC and MS, 8 were DEGs in FE vs FC, and 6 were DEGs in FS vs FC (Figure 4C-D). To investigate the cellular context of these genes, we used the adipose tissue knowledge portal’s^32^ single-cell module with the dataset by Massier et al^33^. Interestingly, all genes in IC 14 were enriched in lipid-associated macrophages (LAM) and metabolically activated macrophages (Figure 4E). Lastly, a transcription factor enrichment analysis (TFEA) indicated that eight of the genes in IC 14 may be regulated by a common transcription factor: activating transcription factor 3 (ATF3). In summary, the ICA detected independent components related to both sex and exercise background. The analysis highlighted a general inflammatory signature of scWAT in untrained individuals, consistent with previous GSEA. Notably, ICA also uncovered a specific set of genes enriched in a macrophage subset, lipid-associated macrophages (LAMs), which showed higher expression in untrained individuals. These findings underscore the utility of ICA for detecting both broad transcriptional patterns and cell type–specific signatures from bulk RNA- seq data.

### Greater abdominal adipose tissue volumes are associated with immune-related pathways

To identify scWAT genes associated with various adipose tissue depot volumes, were correlated gene expression data with abdominal subcutaneous adipose tissue (aSAT) and visceral fat volume determined by a whole-body MRI-scan performed on a subset of the cohort (the MRI data is previously published in^25^). A significant positive correlation between greater aSAT volume and immune system-related GO terms was identified. Specifically, antigen processing and presentation, myeloid leukocyte activation and cell activation involved in immune response were all among the top 10 GO terms (Figure 5A). GO-terms negatively correlated with aSAT volume were primarily related to the ribosome, specifically the cytosolic ribosome, co- translational protein targeting to membrane and large ribosomal subunit but also included fatty acid beta oxidation and mitochondrial transcription (Figure 5A). Seven genes were significantly correlated with aSAT volume (correlation coefficient range 0.82 to 0.74), with γ-synuclein (*SNCG*) displaying the strongest correlation. Next, correlation between gene expression data and visceral fat volume revealed that the top positively correlated GO terms were also immune system-related (Figure 5B). While the most negatively correlated terms in aSAT were related to the ribosome, the terms most negatively correlated with visceral fat volume were all related to aerobic energy production (Figure 5B).

**Figure 5.**
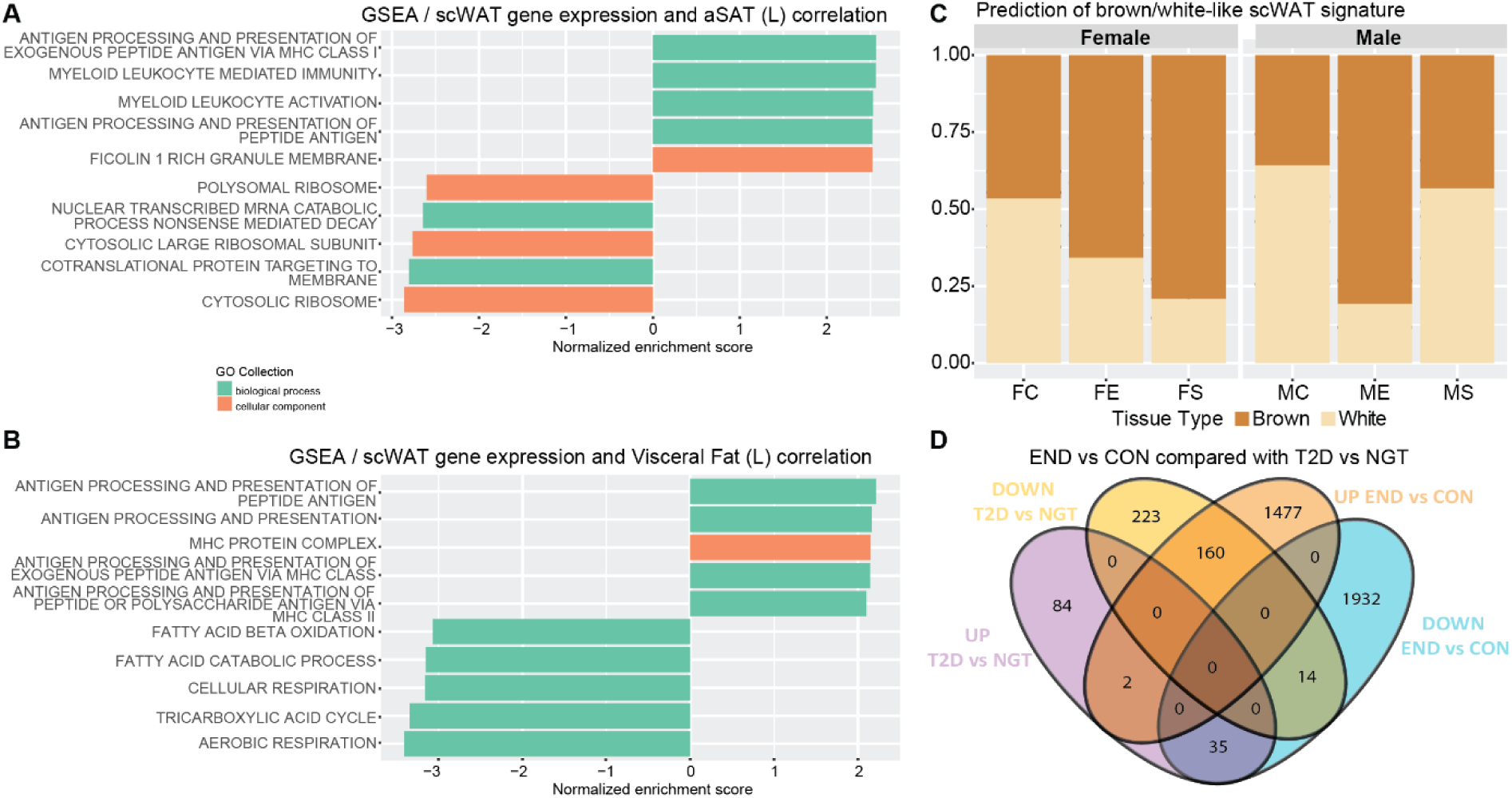
Linking gene expression data with clinical data, scWAT browning and gene expression in T2D patient data. A-B) Gene-set enrichment analysis (GSEA) was performed based on the ranked correlation value following correlations between subcutaneous white adipose tissue (scWAT) RNAseq data and abdominal subcutaneous adipose tissue (aSAT) volume and visceral fat volume, respectively. C) Prediction of the brown- and white-like signature of the scWAT. D) Venn diagram of the up- and downregulated genes in the pooled (males and females) endurance (END) and control (CON) groups of the current study and in a publicly available dataset^18^ comparing gene expression of patients with type 2 diabetes mellitus (T2D) and individuals with normal glucose tolerance (NGT). F = female, M = male, C = control, E = endurance, S = strength, GO = Gene Ontology.

### Long-term endurance trained individuals have a brown-like scWAT signature

In order to further understand the functional remodeling of scWAT with regular exercise training, we analyzed our transcriptomic data with ProFAT^34^ to evaluate the relative brown- and white-like molecular signatures within groups. The highest brown-like signature and thus, the highest thermogenic potential of the scWAT was found in the ME group, followed by FS and FE which all displayed a stronger brown-like than white-like scWAT signature (Figure 5C). The MC displayed the lowest thermogenic potential followed by MS and FC which had a higher thermogenic potential. However, all these groups displayed a more white-like signature (Figure 5C).

### Genes involved in aerobic energy metabolism are oppositely regulated in endurance trained individuals and in patients with type 2 diabetes

To further understand the health benefits of the adaptation to long-term training, we cross-referenced our data with publicly available adipose tissue RNAseq data including male and female patients (however, no sex-specific comparisons were performed in the study^18^) with type 2 diabetes (T2D) and controls with normal glucose tolerance (NGT)^18^. Of the 518 DEGs found comparing T2D with NGT at baseline, 211 were also identified as DEGs in the current study comparing ME vs MC and/or in FE with FC (Figure 5D). An overrepresentation analysis of these 211 genes showed a significant enrichment of the oxidative phosphorylation, thermogenesis and TCA cycle pathways. Additionally, of the 211 genes shared between the respective comparisons, 195 genes were oppositely regulated (160 up in endurance-trained and down in T2D vs NGT), and 16 genes were regulated in the same direction (2 up in both endurance-trained and T2D vs NGT) in endurance trained individuals and in patients with T2D. Among the 195 genes with higher expression in endurance trained and lower in patients with T2D, genes encoding subunits in all five mitochondrial complexes, genes encoding 6 out of 8 TCA cycle enzymes, genes involved in branched chain amino acid metabolism and fatty acid degradation and the *SLC2A4* gene (encoding GLUT4) were detected. Genes involved in apelin signaling, closely associated with brown adipose tissue, and cytokine signaling were found among the 35 genes with higher expression in patients with T2D.

## DISCUSSION

The main finding of the white subcutaneous adipose tissue transcriptomics analysis was that long-term trained athletes had a significantly higher expression of mitochondrial genes, and a lower expression of inflammation-related genes compared with healthy, age-matched untrained individuals. When comparing training backgrounds, the largest number of DEGs was observed between endurance-trained and untrained individuals. Interestingly, large differences were also found between strength trained and untrained females but not in the corresponding male comparison. Independent of specific exercise background, in all trained groups there was a consistent enrichment of GO-terms related to ribosomal and mitochondrial structure and metabolism compared with the corresponding untrained groups. An enrichment of ribosomal and mitochondrial terms in adipose tissue with training is in line with previous data following a six-month long endurance training intervention in healthy males^9^ and in a cross-sectional comparison between endurance trained (>2 years of regular training) and untrained overweight or obese individuals^11^. The enrichment of ribosomal terms in trained individuals may reflect increased expression of ribosomal factors, potentially indicating greater adipogenic capacity and contributing to a higher proportion of smaller adipocytes, which is associated with improved metabolic health^35^.

In the present study, many of the identified genes with the largest fold difference and lowest adjusted p-value between the trained and untrained groups have previously also been identified to show a similar pattern when comparing non-obese versus obese individuals. For example, *EGFL6* and *TM4SF19A,* which were among the top five genes with the highest fold difference in both untrained males and females compared with endurance trained, have previously been shown to be significantly upregulated in obese individuals^32^. Although the current control groups were of normal weight, using MRI measurements we have previously shown that they have a higher aSAT volume, higher total body adipose tissue volume relative to body weight and lower lean body tissue than both the endurance- and strength trained groups^26^. Additionally, both *EGFL6* and *TM4SF19A* have been shown to positively correlate with fat cell volume, homeostatic model assessment for insulin resistance (HOMA-IR), circulating levels of insulin and triglycerides and negatively correlate with circulating HDL^32^. In addition to being upregulated in obese and in untrained individuals, *EGFL6* may act as a paracrine/autocrine growth factor in adipose tissue, contributing to obesity development^36^. Similarly, deletion of *TM4SF19A* in male mice led to adipocyte hyperplasia rather than hypertrophy following high- fat feeding^37^. Furthermore, *SNCG* was identified as the strongest positively correlating gene with aSAT volume. Enhanced *SNCG* expression is associated with obesity in humans, and knockdown experiments in mice led to partial protection against weight gain, reduced both WAT and visceral fat accumulation, and increased lipid oxidation and total energy expenditure^38^. This suggests that individuals performing regular training over many years have a lower expression of genes associated with an unhealthy metabolic profile and lower scWAT volume, compared with non-exercisers. On a similar note, we show that scWAT from long-term endurance-trained males and females, as well as strength-trained females, displays a brown-like adipose signature. This aligns with previous findings of an increased expression of brown/beige associated genes following 12 weeks of endurance training^39^. This is likely driven by the higher expression of mitochondrial genes in these exercised-trained groups.

The independent component analysis identified several biologically relevant gene sets related to both training background and sex. Of particular interest is IC 14, as all 11 genes were tightly co-regulated, had a higher expression in untrained than in trained as well as in obese compared with non-obese individuals^32^, and exhibit distinct enrichment in lipid associated macrophages and in metabolically activated macrophages. Two of the genes, *CHI3L1* and *IL1RN*, were also found to be upregulated in patients with T2D compared with those with NGT^18^. Previous research has shown a strong positive correlation between lipid-associated macrophages and BMI in human visceral adipose tissue^40^, and that they are related to detrimental metabolic phenotypes such as impaired lipid mobilization, larger fat cell volume and increased waist-to- hip ratio^33^. Moreover, mouse models have identified LAM cells to be the most expanded immune cell subset in obesity^40^, and their formation is suggested to be driven by TREM2 signaling^40^. *TREM2* expression was significantly lower in all trained groups compared with controls, except in strength-trained versus untrained males. Furthermore, ATF3 was identified as a candidate TF based on co-expression. ATF3 has previously been shown to regulate lipid-loading and lipid metabolic pathways in macrophages^41^. Its mRNA expression is increased in obese individuals^42^, and overexpression of ATF3 in transgenic mice has been shown to reduce the expression of mitochondrial genes^43^. However, ATF3 overexpression, in mice, has also been suggested to induce browning^44^. Thus, the role of ATF3 in adipose tissue remains unclear, and further investigation is warranted, particularly in human scWAT. Interestingly, the vast majority of genes in component 14 were identified as DEGs in the endurance-trained males and females, as well as in strength-trained females, but not in strength-trained males, when compared with their respective control. This suggests a possible sex-specific adaptation in scWAT following strength training. Specifically, it implies that in males, strength training alone may not be sufficient to downregulate genes enriched in LAM cells. This analysis also highlights the utility of ICA as a useful tool for identification of important gene sets. Lastly, given the apparent involvement of these genes in metabolic processes within scWAT, further mechanistic studies are warranted to clarify their role in human metabolism and to explore possible sex-specific adaptations to strength training.

A distinct sexually dimorphic characteristic of scWAT was identified in this study. Specifically, sex differences were largely driven by an enrichment of genes and pathways related to aerobic metabolism in females, and by inflammation-related pathways in males. These findings align with previous studies reporting higher subcutaneous adipose tissue expression of nuclear-encoded OXPHOS genes^45^, enrichment of OXPHOS and fatty acid metabolism pathways in females^46^, as well as sex-specific differences in inflammation-related gene expression^46^. Similar enrichment of aerobic metabolism pathways has also been observed in skeletal muscle from untrained females compared to males^47^. Prior research has demonstrated that the immune system operates differently between the sexes, in part due to the modulatory effects of sex hormones^48^. As such, a recent study showed that testosterone modulates pro-inflammatory signaling via the type-I interferon and tumor necrosis factor axis^48^, which partially can explain the pro-inflammatory transcriptome observed in male scWAT. Supporting this notion, another study found that knockout of estrogen receptor-α in adipose tissue led to pro-fibrotic and pro-inflammatory effects in a mouse model^49^, further highlighting the role of sex hormones in regulating adipose tissue inflammation. Taken together, these findings support the existence of a more pro-inflammatory scWAT transcriptomic profile in males compared with females. However, our data also suggest that regular physical training attenuates this inflammatory profile of male scWAT. Notably, endurance training appears to offer additional benefits, as no immune system-related terms were significantly enriched in endurance-trained males compared with females. Therefore, to fully counteract the inherent pro-inflammatory transcriptional profile in male scWAT, it may be necessary to incorporate endurance exercise into training routines. Thus, this sexual dimorphism may reflect evolutionary pressures whereby premenopausal females evolved to maintain insulin-sensitive scWAT^50^ to support reproductive and energetic demands, whereas males evolved a more pro-inflammatory profile that, while potentially advantageous for acute survival, compromises metabolic flexibility and predisposes to long-term systemic consequences Additionally, many of the top DEGs in females versus males, such as HACD2 (up in FE vs ME), ABCC9 (up in FE vs ME), and MESP1 (up in FS vs MS), have previously been shown to be more highly expressed in non-obese versus obese individuals and negatively correlate with HOMA-IR, fat cell volume, and circulating triglycerides^32^. In contrast, several top genes in males show higher expression in obese versus non-obese individuals and correlate positively with HOMA-IR and triglycerides but negatively with HDL^32^. Taken together, these findings suggest that females have an inherently metabolically beneficial scWAT gene expression profile, which likely explains the larger differences observed between endurance-trained males and their controls relative to females.

In conclusion, our study demonstrates that long-term highly trained individuals have a significantly altered scWAT transcriptome compared with age-matched healthy untrained controls. The differences included an altered expression of genes that have previously been linked to metabolic diseases such as obesity and type 2 diabetes, suggesting a protective effect of long-term training, particularly through endurance training, and to a greater extent in strength-trained females than in males. Notably, we identified a set of 11 co-regulated genes enriched in lipid-associated macrophages that were co-regulated in scWAT. Interestingly, eight of these genes were predicted to be regulated by the transcription factor ATF3, suggesting a potential upstream regulatory mechanism. This gene set exhibited opposing regulation patterns between endurance-trained males and females, as well as between strength-trained females and obese individuals—an effect not observed in strength-trained males. This finding suggests that, in males, endurance training may be necessary to fully counteract the enrichment of inflammatory pathways in scWAT. Future studies should target this gene set further to examine its role in scWAT metabolism in relation to exercise training. The inflammatory profile of the male compared with female scWAT was attenuated with long-term training, yet to a greater extent in endurance trained males. These results underscore the sexually dimorphic nature of the scWAT transcriptome and emphasize the importance of including both sexes in adipose tissue research. Sex-specific analyses will ultimately enable the development of more precise diagnostic tools and targeted treatments in metabolic health.

## Acknowledgements

This work was financially supported by grants from the Swedish Research Council (2018-02932), the Swedish Center for Sports Research (D2018-0007, P2022-0082), the National Science Foundation (Award Number: 1951792), the Whitaker International Program, and Karolinska Institutet institutional funding. The authors acknowledge support from the National Genomics Infrastructure in Stockholm funded by Science for Life Laboratory, the Knut and Alice Wallenberg Foundation and the Swedish Research Council, and NAISS/Uppsala Multidisciplinary Center for Advanced Computational Science for assistance with massively parallel sequencing and access to the UPPMAX computational infrastructure. Figure 1A was created with BioRender.com.

## Author contributions

Conceptualization, E.B.E., S.M.R., D.C.Z, M.R., M.A.C., and C.J.S.,; methodology, E.B.E., S.M.R., D.C.Z, M.R., M.A.C., and C.J.S.; formal analysis, E.B.E., S.M.R., A.Q., J.T.B., and D.C.Z.; investigation, E.B.E, S.M.R., A.Q., J.T.B., J.J., M.E., H.G., S.P., M.A.C., and C.J.S.; project management C.J.S.; resources B.O.P., M.R., and C.J.S.; visualization, E.B.E, S.M.R., A.Q., J.T.B., and J.J.; Supervision, M.E.L., J.N., M.R, B.O.P., D.C.Z., M.A.C., and C.J.S.; funding acquisition M.A.C., and C.J.S.; writing—original draft, E.B.E., and S.M.R.; writing— review & editing, all authors.

## Competing Interests

The authors declare no competing interests.

## Methods

### Ethics statement

This research received approval from the Regional Ethical Review Board in Stockholm, Sweden (application 2016/590-31) and adhered to the Declaration of Helsinki. Before the experiment began, each participant was given a comprehensive explanation of all procedures and possible risks. Written and verbal consent were obtained, and participants were informed that they could withdraw their consent at any point during the study.

### Subjects and experimental design

A total of 89 individuals (47 females and 42 males) were included into three separate groups based on exercise training background, an untrained control group, a highly endurance trained group and a highly strength trained group. The inclusion of participants has been described previously^47^. In short, the endurance-trained males (ME) and females (FE) had a significantly higher VO_2-peak,_ by design, than both respective control and strength-trained groups (Figure 1C). The strength-trained males (MS) and females (FS) were significantly stronger than the respective control and endurance groups (Figure 1C). The male and female control groups (MC and FC, respectively) consisted of healthy age-matched, untrained individuals with no chronic conditions or medications and a BMI ≤ 27. All trained individuals have at least 15 years’ experience of regular training in their respective sports. A subset (n = 64) of the cohort has already been included in previous studies focusing on the body composition and molecular adaptations in skeletal muscle of the same individuals^26,27,47,52^.

### Biopsy collection

Paraumbilical scWAT biopsies were taken at rest following local anesthesia with aspiration of Hepafix liver biopsy kits (B. Braun, Germany). After the adipose tissue biopsies were washed with NaCl and cleared of capillaries, the tissue was snap-frozen in liquid nitrogen. All biopsies were collected following a standardized breakfast, at least 72 hours after the last exercise bout as previously described^47^.

### Histological staining and analysis

Adipocyte area was quantified on a subset of the cohort (n = 18, three individuals from each group). Frozen adipose tissue samples were embedded in optimal cutting temperature compound, cut in 7 µm thick sections and mounted onto glass slides. First, the slides were dipped into 4% PFA for 30 seconds. They were then immersed in hematoxylin (Mayers Hematoxylin Plus #01825) for 4 minutes, followed by washing under running water for 5 minutes. Next, the slides were dipped into eosin (ready-made 0.2% solution #01650) for 3 minutes and briefly washed in water. For dehydration, the slides underwent a series of ethanol baths. In the final step, the slides were dipped in xylene for 30 seconds and then covered with xylene-based mounting media^53^. Images were acquired using a 3D HISTECH PANNORAMIC MIDI II scanner (RRID:SCR_024834) with a 40x objective and a and Hitachi (HV-F22CL) color camera. Cell area was quantified manually by a single investigator, who was completely blinded to the experimental groups and sex, using imageJ (ImageJ, RRID:SCR_003070). Due to the low sample size (n = 3 per group) no statistical analyses were performed on the adipocyte sizes and proportions.

#### RNA sequencing

RNA extraction from the adipose tissue was performed using the phenol based TRIzol method (Invitrogen #15596018, Thermo Fisher Scientific, USA). Libraries were prepared by poly-A selection (TruSeq mRNA, Illumina, San Diego, CA, USA) and multiplexed at the National Genomics Infrastructure Sweden. Clustering was done by ’cBot’ and samples were sequenced on NovaSeq6000 (NovaSeq Control Software 1.6.0/RTA v3.4.4) with two lanes of a 2x151 setup ’NovaSeqXp’ workflow in ’S4’ mode flow cell. The Bcl to FastQ conversion was performed using bcl2fastq_v2.20.0.422 from the CASAVA software suite. The quality scale used is Sanger / phred33 / Illumina 1.8+. QC and processing were performed using the nfcore/rnaseq analysis pipeline publicly available at github (https://github.com/nf-core/rnaseq). *Bioinformatics*

To perform statistical analyses on the RNA sequencing data, genes were first selected based on expression thresholds of >0.1 TPM in at least 20% of the samples and ≥6 reads in at least 20% of the samples. This selection criteria comes from the GTEx Portal^54^. Statistical analyses were then conducted with R, using the raw counts as the input to DESeq2^55^ to perform principal component analysis and differential gene expression analysis, with a false discovery rate (FDR) set to 5%. Gene set enrichment analysis (GSEA) was performed with the fGSEA package^56^ using Gene Ontology terms (biological process, BP; molecular function, MF; and cellular components, CC)^57,58^. For comparison of mitochondria-specific pathways, the GSEA was performed using MitoPathways 3.0^23^ using the fGSEA package^56^. CIBERSORTx was used to infer the immune cell type proportions between groups and sexes using the curated signature matrix LM22 signature matrix, as described elsewhere^28^. The CIBERSORTx run was performed following bulk mode batch correction, run in absolute mode with 100 permutations. Correlation analysis of the MRI and gene expression were performed using the Pearson correlation coefficient on all individuals where we had both MRI and RNAseq data (n = 32). It should be noted that the FS groups were missing in this analysis. GSEA was subsequently performed based on a ranking by correlation value r. Overrepresentation analysis was performed with Enrichr^59^ with the genes included in the differential gene expression analysis as background data. All GSEA and overrepresentation analyses defined significant pathways/GO terms as FDR ≤0.05. Prediction of brown-like signature was performed using ProFAT^34^. Comparison of the identified DEGs in the current study with the DEGs from Dollet et al^18^ were made through directional comparisons, and genes were matched by gene symbols. The transcription factor enrichment analysis was performed using the ChIP-X Enrichment Analysis Version 3 (ChEA3)^60^ database with the ARCHS4 dataset as selected library.

#### Independent Component Analysis

Independent component analysis was performed in order to decompose the expression of genes across RNA-seq profiles into two matrices, one of independent components (iModulons) in the M matrix and one of their activities across profiles, the A matrix^61^. Feature selection was performed prior to model execution by filtering genes with mean counts fewer than 1 per sample and then those remaining genes which were not in the top 70% of variation across all genes, resulting in 10343 genes being included in the model. The counts were normalized using DESeq2^55^. Prior to running ICA, the mean expression of genes in MC was subtracted from all samples to provide a point of comparison. The iModulon Miner workflow was used to perform ICA across ranks 10 to 80 in steps of 10, with each rank being performed 100 times and the robust clusters determined through clustering with DBSCAN^62^ with an epsilon of .1 and a minimum cluster seed size of 50. Rank 60 was selected for further analysis based on established protocols^63^. Finally, the highest weight gene in each iModulon was set to be positive and the associated genes and activities were adjusted accordingly. The gene membership of each iModulon was achieved using 3-means clustering^64^ and the two clusters with the highest absolute value gene weights were averaged in order to set the threshold value for gene membership in an iModulon.

#### Statistical analysis

Subject characteristics were calculated using two-way ANOVAs, with Tukey’s post hoc test used in case of statistically significant group differences from the ANOVAs. Statistical analyses were performed using GraphPad Prism 10 (GraphPad, Prism, RRID:SCR_002798). scWAT transcriptomic analysis was performed as previously described, see section “bioinformatics”.

## Data and Code Availability

The raw sequencing data generated in this study is deposited in the European Genome-phenome Archive (EGA) and will be publicly available following publication of the study. Scripts used for the analyses in the study will be available at GitHub following publication.

## Supplementary Data

**Supplemental Data 1.** List of differentially expressed genes, fGSEA results using GO terms, and fGSEA results based on MitoPathways.

**Supplemental Data 2.**
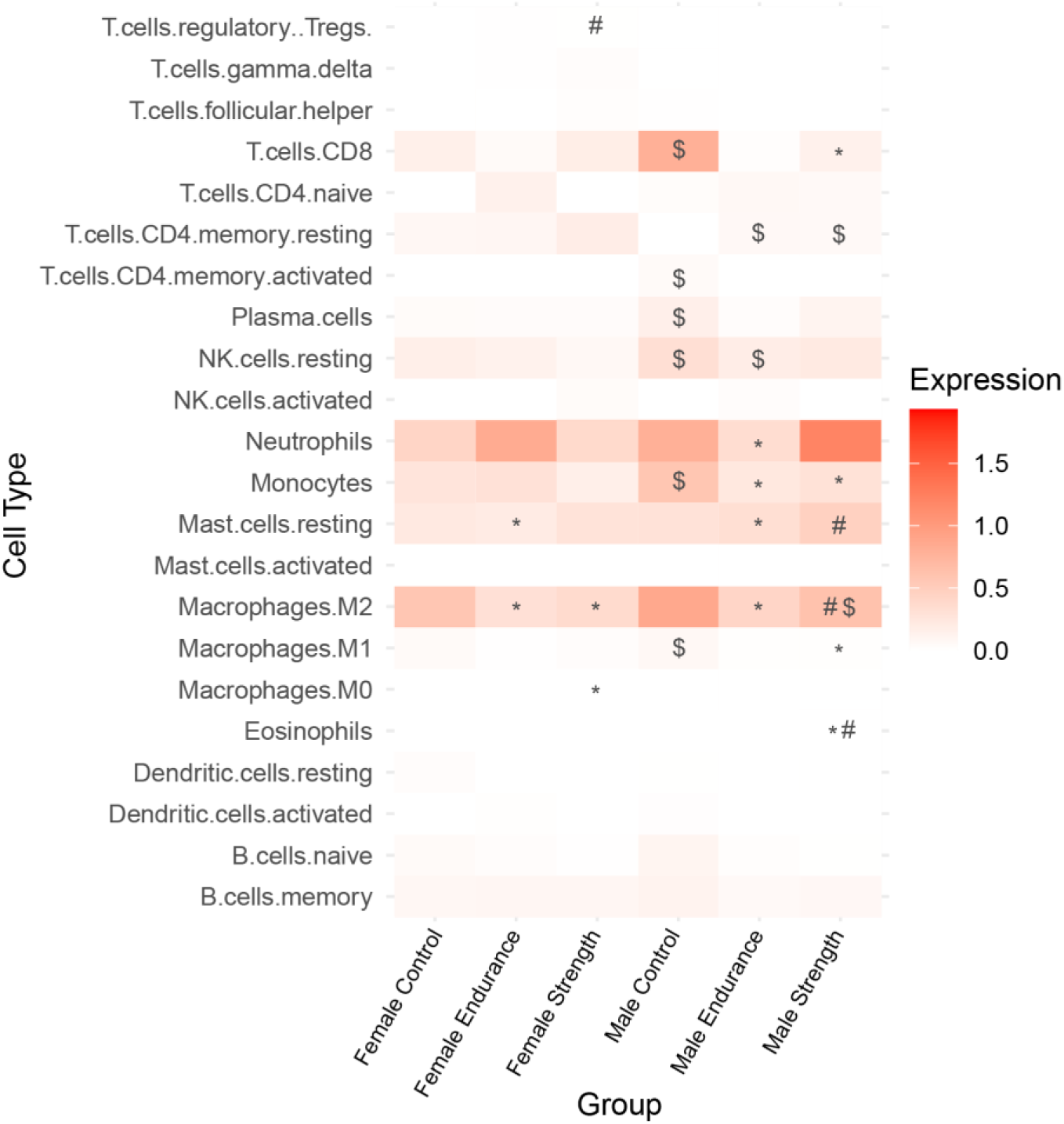
CIBERSORTx analysis. * = Significantly different to sex-matched control group, # = Significantly different to sex-matched endurance group, $ = Significant sex difference (e.g. male control vs female control).

**Supplemental data 3.**
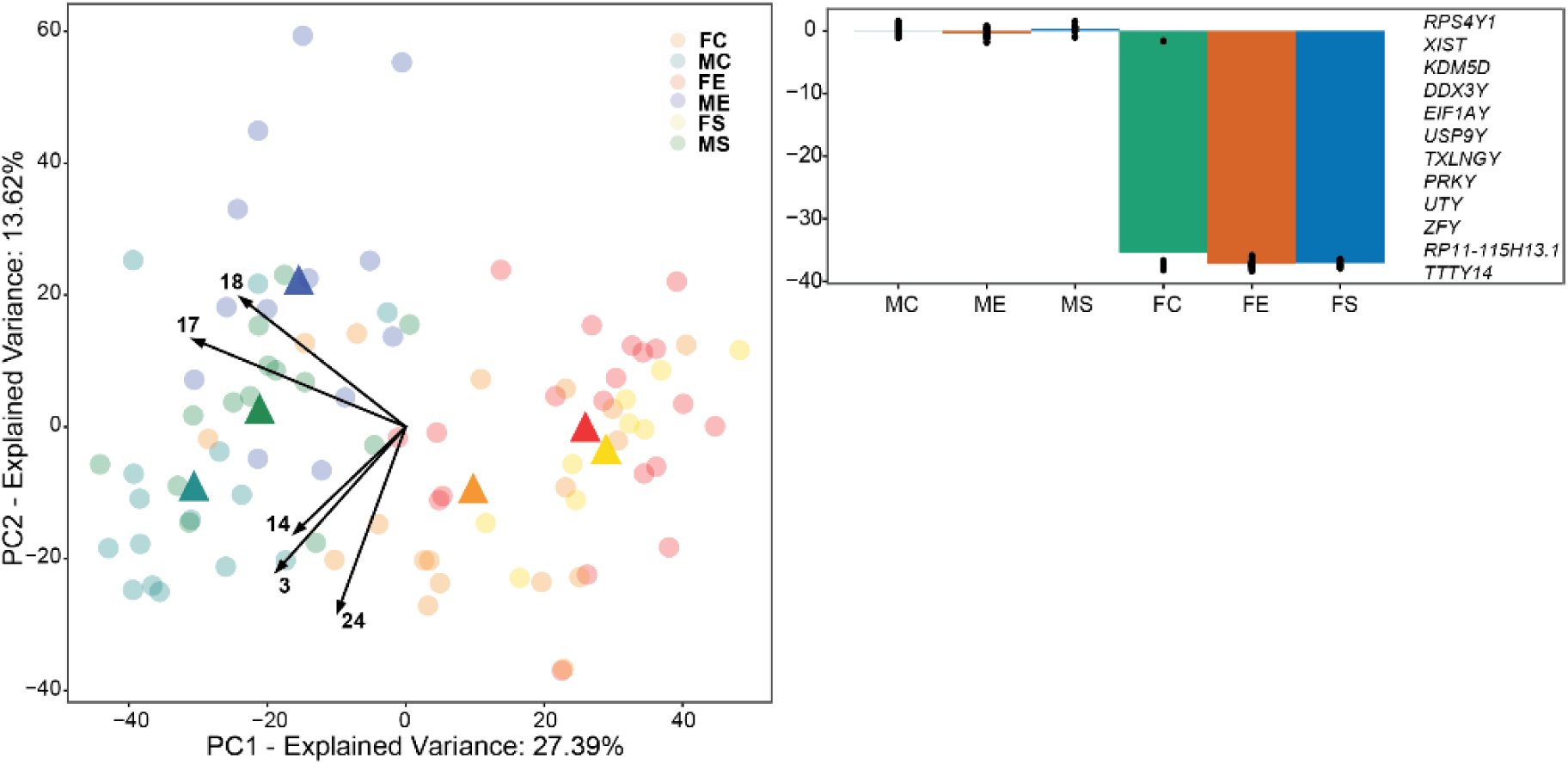
Independent component analysis. A) PCA biplot of the ICA data. Circles represent each individual; triangles display the centroids for each group; the arrows represent the direction of the top five independent components (ICs) with the largest explained variance. B) Displays the activity of the Y-linked IC 17, with the average activity for each group and a list of all genes included in the IC.

## Notes

### Competing Interest Statement

The authors have declared no competing interest.

## References

1. Zhang, Y., Proenca, R., Maffei, M., Barone, M., Leopold, L., and Friedman, J.M. (1994). Positional cloning of the mouse obese gene and its human homologue. Nature 372, 425–432. 10.1038/372425a0.

2. Hotamisligil, G.S., Shargill, N.S., and Spiegelman, B.M. (1993). Adipose expression of tumor necrosis factor-alpha: direct role in obesity-linked insulin resistance. Science 259, 87–91. 10.1126/science.7678183.

3. Després, J.P., Bouchard, C., Savard, R., Tremblay, A., Marcotte, M., and Thériault, G. (1984). The effect of a 20-week endurance training program on adipose-tissue morphology and lipolysis in men and women. Metabolism. 33, 235–239. 10.1016/0026-0495(84)90043-x.

4. Stallknecht, B., Larsen, J.J., Mikines, K.J., Simonsen, L., Bülow, J., and Galbo, H. (2000). Effect of training on insulin sensitivity of glucose uptake and lipolysis in human adipose tissue. Am. J. Physiol.-Endocrinol. Metab. 279, E376–E385. 10.1152/ajpendo.2000.279.2.E376.

5. Moore, T.M., Lee, S., Olsen, T., Morselli, M., Strumwasser, A.R., Lin, A.J., Zhou, Z., Abrishami, A., Garcia, S.M., Bribiesca, J., et al. (2023). Conserved multi-tissue transcriptomic adaptations to exercise training in humans and mice. Cell Rep. 42, 112499. 10.1016/j.celrep.2023.112499.

6. Horowitz, J.F. (2003). Fatty acid mobilization from adipose tissue during exercise. Trends Endocrinol. Metab. 14, 386–392. 10.1016/S1043-2760(03)00143-7.

7. Walton, R.G., Finlin, B.S., Mula, J., Long, D.E., Zhu, B., Fry, C.S., Westgate, P.M., Lee, J.D., Bennett, T., Kern, P.A., et al. (2015). Insulin-resistant subjects have normal angiogenic response to aerobic exercise training in skeletal muscle, but not in adipose tissue. Physiol. Rep. 3, e12415. 10.14814/phy2.12415.

8. Riis, S., Christensen, B., Nellemann, B., Møller, A.B., Husted, A.S., Pedersen, S.B., Schwartz, T.W., Jørgensen, J.O.L., and Jessen, N. (2019). Molecular adaptations in human subcutaneous adipose tissue after ten weeks of endurance exercise training in healthy males. J. Appl. Physiol. Bethesda Md 1985 *126*, 569–577. 10.1152/japplphysiol.00989.2018.

9. Rönn, T., Volkov, P., Tornberg, A., Elgzyri, T., Hansson, O., Eriksson, K.-F., Groop, L., and Ling, C. (2014). Extensive changes in the transcriptional profile of human adipose tissue including genes involved in oxidative phosphorylation after a 6-month exercise intervention. Acta Physiol. Oxf. Engl. 211, 188–200. 10.1111/apha.12247.

10. Pino, M.F., Dijkstra, P., Whytock, K.L., Ahn, C., Yu, G., Sanford, J.A., Hansen, J., Hutchinson, C., Gritsenko, M., Piehowski, P., et al. (2025). Exercise alters molecular profiles of inflammation and substrate metabolism in human white adipose tissue. Am. J. Physiol. Endocrinol. Metab. 328, E478–E492. 10.1152/ajpendo.00339.2024.

11. Ahn, C., Zhang, T., Yang, G., Rode, T., Varshney, P., Ghayur, S.J., Chugh, O.K., Jiang, H., and Horowitz, J.F. (2024). Years of endurance exercise training remodel abdominal subcutaneous adipose tissue in adults with overweight or obesity. Nat. Metab. 6, 1819– 1836. 10.1038/s42255-024-01103-x.

12. Gudiksen, A., Qoqaj, A., Ringholm, S., Wojtaszewski, J., Plomgaard, P., and Pilegaard, H. (2022). Ameliorating Effects of Lifelong Physical Activity on Healthy Aging and Mitochondrial Function in Human White Adipose Tissue. J. Gerontol. Ser. A 77, 1101–1111. 10.1093/gerona/glab356.

13. Stinkens, R., Brouwers, B., Jocken, J.W., Blaak, E.E., Teunissen-Beekman, K.F., Hesselink, M.K., van Baak, M.A., Schrauwen, P., and Goossens, G.H. (2018). Exercise training-induced effects on the abdominal subcutaneous adipose tissue phenotype in humans with obesity. J. Appl. Physiol. Bethesda Md 1985 *125*, 1585–1593. 10.1152/japplphysiol.00496.2018.

14. Kang, S., Park, K.-M., Sung, K.-Y., Yuan, Y., and Lim, S.-T. (2021). Effect of Resistance Exercise on the Lipolysis Pathway in Obese Pre- and Postmenopausal Women. J. Pers. Med. 11, 874. 10.3390/jpm11090874.

15. Klimcakova, E., Polak, J., Moro, C., Hejnova, J., Majercik, M., Viguerie, N., Berlan, M., Langin, D., and Stich, V. (2006). Dynamic strength training improves insulin sensitivity without altering plasma levels and gene expression of adipokines in subcutaneous adipose tissue in obese men. J. Clin. Endocrinol. Metab. 91, 5107–5112. 10.1210/jc.2006-0382.

16. Fabre, O., Ingerslev, L.R., Garde, C., Donkin, I., Simar, D., and Barrès, R. (2018). Exercise training alters the genomic response to acute exercise in human adipose tissue. Epigenomics 10, 1033–1050. 10.2217/epi-2018-0039.

17. Van Pelt, D.W., Guth, L.M., and Horowitz, J.F. (2017). Aerobic exercise elevates markers of angiogenesis and macrophage IL-6 gene expression in the subcutaneous adipose tissue of overweight-to-obese adults. J. Appl. Physiol. 123, 1150–1159. 10.1152/japplphysiol.00614.2017.

18. Dollet, L., Lundell, L.S., Chibalin, A.V., Pendergrast, L.A., Pillon, N.J., Lansbury, E.L., Elmastas, M., Frendo-Cumbo, S., Jalkanen, J., de Castro Barbosa, T., et al. (2023). Exercise-induced crosstalk between immune cells and adipocytes in humans: Role of oncostatin-M. Cell Rep. Med. 5, 101348. 10.1016/j.xcrm.2023.101348.

19. Gershoni, M., and Pietrokovski, S. (2017). The landscape of sex-differential transcriptome and its consequent selection in human adults. BMC Biol. 15, 7. 10.1186/s12915-017-0352-z.

20. Oliva, M., Muñoz-Aguirre, M., Kim-Hellmuth, S., Wucher, V., Gewirtz, A.D.H., Cotter, D.J., Parsana, P., Kasela, S., Balliu, B., Viñuela, A., et al. (2020). The impact of sex on gene expression across human tissues. Science 369, eaba3066. 10.1126/science.aba3066.

21. Chang, E., Varghese, M., and Singer, K. (2018). Gender and Sex Differences in Adipose Tissue. Curr. Diab. Rep. 18, 69. 10.1007/s11892-018-1031-3.

22. Many, G.M., Sanford, J.A., Sagendorf, T.J., Hou, Z., Nigro, P., Whytock, K.L., Amar, D., Caputo, T., Gay, N.R., Gaul, D.A., et al. (2024). Sexual dimorphism and the multi-omic response to exercise training in rat subcutaneous white adipose tissue. Nat. Metab. 6, 963–979. 10.1038/s42255-023-00959-9.

23. Rath, S., Sharma, R., Gupta, R., Ast, T., Chan, C., Durham, T.J., Goodman, R.P., Grabarek, Z., Haas, M.E., Hung, W.H.W., et al. (2021). MitoCarta3.0: an updated mitochondrial proteome now with sub-organelle localization and pathway annotations. Nucleic Acids Res. 49, D1541–D1547. 10.1093/nar/gkaa1011.

24. Saltin, B., and Astrand, P.O. (1967). Maximal oxygen uptake in athletes. J. Appl. Physiol. 23, 353–358. 10.1152/jappl.1967.23.3.353.

25. Häkkinen, K., and Keskinen, K.L. (1989). Muscle cross-sectional area and voluntary force production characteristics in elite strength- and endurance-trained athletes and sprinters. Eur. J. Appl. Physiol. 59, 215–220.

26. Emanuelsson, E.B., Berry, D.B., Reitzner, S.M., Arif, M., Mardinoglu, A., Gustafsson, T., Ward, S.R., Sundberg, C.J., and Chapman, M.A. (2022). MRI characterization of skeletal muscle size and fatty infiltration in long-term trained and untrained individuals. Physiol. Rep. 10, e15398. 10.14814/phy2.15398.

27. Reitzner, S.M., Emanuelsson, E.B., Arif, M., Kaczkowski, B., Kwon, A.TJ., Mardinoglu, A., Arner, E., Chapman, M.A., and Sundberg, C.J. (2024). Molecular profiling of high-level athlete skeletal muscle after acute endurance or resistance exercise – A systems biology approach. Mol. Metab. 79, 101857. 10.1016/j.molmet.2023.101857.

28. Newman, A.M., Steen, C.B., Liu, C.L., Gentles, A.J., Chaudhuri, A.A., Scherer, F., Khodadoust, M.S., Esfahani, M.S., Luca, B.A., Steiner, D., et al. (2019). Determining cell type abundance and expression from bulk tissues with digital cytometry. Nat. Biotechnol. 37, 773–782. 10.1038/s41587-019-0114-2.

29. Karczewski, K.J., Snyder, M., Altman, R.B., and Tatonetti, N.P. (2014). Coherent Functional Modules Improve Transcription Factor Target Identification, Cooperativity Prediction, and Disease Association. PLOS Genet. 10, e1004122. 10.1371/journal.pgen.1004122.

30. Engreitz, J.M., Daigle, B.J., Marshall, J.J., and Altman, R.B. (2010). Independent component analysis: mining microarray data for fundamental human gene expression modules. J.Biomed. Inform. 43, 932–944. 10.1016/j.jbi.2010.07.001.

31. Rychel, K., Decker, K., Sastry, A.V., Phaneuf, P.V., Poudel, S., and Palsson, B.O. (2020). iModulonDB: a knowledgebase of microbial transcriptional regulation derived from machine learning. Nucleic Acids Res. 49, D112–D120. 10.1093/nar/gkaa810.

32. Zhong, J., Zareifi, D., Weinbrenner, S., Hansen, M., Klingelhuber, F., Nono Nankam, P.A., Frendo-Cumbo, S., Bhalla, N., Cordeddu, L., de Castro Barbosa, T., et al. (2025). adiposetissue.org: A knowledge portal integrating clinical and experimental data from human adipose tissue. Cell Metab., S1550–4131(25)00012-9. 10.1016/j.cmet.2025.01.012.

33. Massier, L., Jalkanen, J., Elmastas, M., Zhong, J., Wang, T., Nono Nankam, P.A., Frendo-Cumbo, S., Bäckdahl, J., Subramanian, N., Sekine, T., et al. (2023). An integrated single cell and spatial transcriptomic map of human white adipose tissue. Nat. Commun. 14, 1438. 10.1038/s41467-023-36983-2.

34. Cheng, Y., Jiang, L., Keipert, S., Zhang, S., Hauser, A., Graf, E., Strom, T., Tschöp, M., Jastroch, M., and Perocchi, F. (2018). Prediction of Adipose Browning Capacity by Systematic Integration of Transcriptional Profiles. Cell Rep. 23, 3112–3125. 10.1016/j.celrep.2018.05.021.

35. De Siqueira, M.K., Li, G., Zhao, Y., Wang, S., Ahn, I.S., Tamboline, M., Hildreth, A.D., Larios, J., Schcolnik-Cabrera, A., Nouhi, Z., et al. (2024). PPARγ-dependent remodeling of translational machinery in adipose progenitors is impaired in obesity. Cell Rep. 43, 114945. 10.1016/j.celrep.2024.114945.

36. Oberauer, R., Rist, W., Lenter, M.C., Hamilton, B.S., and Neubauer, H. (2010). EGFL6 is increasingly expressed in human obesity and promotes proliferation of adipose tissue-derived stromal vascular cells. Mol. Cell. Biochem. 343, 257–269. 10.1007/s11010-010-0521-7.

37. Choi, C., Jeong, Y.L., Park, K.-M., Kim, M., Kim, S., Jo, H., Lee, S., Kim, H., Choi, G., Choi, Y.H., et al. (2024). TM4SF19-mediated control of lysosomal activity in macrophages contributes to obesity-induced inflammation and metabolic dysfunction. Nat. Commun. 15, 2779. 10.1038/s41467-024-47108-8.

38. Millership, S., Ninkina, N., Guschina, I.A., Norton, J., Brambilla, R., Oort, P.J., Adams, S.H., Dennis, R.J., Voshol, P.J., Rochford, J.J., et al. (2012). Increased lipolysis and altered lipid homeostasis protect γ-synuclein–null mutant mice from diet-induced obesity. Proc. Natl. Acad. Sci. 109, 20943–20948. 10.1073/pnas.1210022110.

39. Otero-Díaz, B., Rodríguez-Flores, M., Sánchez-Muñoz, V., Monraz-Preciado, F., Ordoñez-Ortega, S., Becerril-Elias, V., Baay-Guzmán, G., Obando-Monge, R., García-García, E., Palacios-González, B., et al. (2018). Exercise Induces White Adipose Tissue Browning Across the Weight Spectrum in Humans. Front. Physiol. 9. 10.3389/fphys.2018.01781.

40. Jaitin, D.A., Adlung, L., Thaiss, C.A., Weiner, A., Li, B., Descamps, H., Lundgren, P., Bleriot, C., Liu, Z., Deczkowska, A., et al. (2019). Lipid-associated macrophages control metabolic homeostasis in a Trem2-dependent manner. Cell 178, 686–698.e14. 10.1016/j.cell.2019.05.054.

41. Gold, E.S., Ramsey, S.A., Sartain, M.J., Selinummi, J., Podolsky, I., Rodriguez, D.J., Moritz, R.L., and Aderem, A. (2012). ATF3 protects against atherosclerosis by suppressing 25-hydroxycholesterol–induced lipid body formation. J. Exp. Med. 209, 807–817. 10.1084/jem.20111202.

42. Hu, S., Cassim Bawa, F.N., Zhu, Y., Pan, X., Wang, H., Gopoju, R., Xu, Y., and Zhang, Y. (2024). Loss of adipose ATF3 promotes adipose tissue lipolysis and the development of MASH. Commun. Biol. 7, 1–12. 10.1038/s42003-024-06915-x.

43. Jang, M.-K., Son, Y., and Jung, M.H. (2013). ATF3 plays a role in adipocyte hypoxia-mediated mitochondria dysfunction in obesity. Biochem. Biophys. Res. Commun. 431, 421–427. 10.1016/j.bbrc.2012.12.154.

44. Cheng, C.-F., Ku, H.-C., Cheng, J.-J., Chao, S.-W., Li, H.-F., Lai, P.-F., Chang, C.-C., Don, M.-J., Chen, H.-H., and Lin, H. (2019). Adipocyte browning and resistance to obesity in mice is induced by expression of ATF3. Commun. Biol. 2, 389. 10.1038/s42003-019-0624-y.

45. Chella Krishnan, K., Vergnes, L., Acín-Pérez, R., Stiles, L., Shum, M., Ma, L., Mouisel, E., Pan, C., Moore, T.M., Péterfy, M., et al. (2021). Sex-specific genetic regulation of adipose mitochondria and metabolic syndrome by Ndufv2. Nat. Metab. 3, 1552–1568. 10.1038/s42255-021-00481-w.

46. Anderson, W.D., Soh, J.Y., Innis, S.E., Dimanche, A., Ma, L., Langefeld, C.D., Comeau, M.E., Das, S.K., Schadt, E.E., Björkegren, J.L.M., et al. (2020). Sex differences in human adipose tissue gene expression and genetic regulation involve adipogenesis. Genome Res. 30, 1379– 1392. 10.1101/gr.264614.120.

47. Chapman, M.A., Arif, M., Emanuelsson, E.B., Reitzner, S.M., Lindholm, M.E., Mardinoglu, A., and Sundberg, C.J. (2020). Skeletal Muscle Transcriptomic Comparison between Long-Term Trained and Untrained Men and Women. Cell Rep. 31, 107808. 10.1016/j.celrep.2020.107808.

48. Lakshmikanth, T., Consiglio, C., Sardh, F., Forlin, R., Wang, J., Tan, Z., Barcenilla, H., Rodriguez, L., Sugrue, J., Noori, P., et al. (2024). Immune system adaptation during gender-affirming testosterone treatment. Nature 633, 155–164. 10.1038/s41586-024-07789-z.

49. Davis, K.E., D Neinast, M., Sun, K., M Skiles, W., D Bills, J., A Zehr, J., Zeve, D., D Hahner, L., W Cox, D., M Gent, L., et al. (2013). The sexually dimorphic role of adipose and adipocyte estrogen receptors in modulating adipose tissue expansion, inflammation, and fibrosis. Mol. Metab. 2, 227–242. 10.1016/j.molmet.2013.05.006.

50. Mauvais-Jarvis, F. (2015). Sex differences in metabolic homeostasis, diabetes, and obesity. Biol. Sex Differ. 6, 14. 10.1186/s13293-015-0033-y.

51. Wang, H., and Ye, J. (2015). Regulation of energy balance by inflammation: common theme in physiology and pathology. Rev. Endocr. Metab. Disord. 16, 47–54. 10.1007/s11154-014-9306-8.

52. Emanuelsson, E.B., Arif, M., Reitzner, S.M., Perez, S., Lindholm, M.E., Mardinoglu, A., Daub, C., Sundberg, C.J., and Chapman, M.A. (2024). Remodeling of the human skeletal muscle proteome found after long-term endurance training but not after strength training. iScience 27. 10.1016/j.isci.2023.108638.

53. Parlee, S.D., Lentz, S.I., Mori, H., and MacDougald, O.A. (2014). Quantifying Size and Number of Adipocytes in Adipose Tissue. Methods Enzymol. 537, 93–122. 10.1016/B978-0-12-411619-1.00006-9.

54. Lonsdale, J., Thomas, J., Salvatore, M., Phillips, R., Lo, E., Shad, S., Hasz, R., Walters, G., Garcia, F., Young, N., et al. (2013). The Genotype-Tissue Expression (GTEx) project. Nat. Genet. 45, 580–585. 10.1038/ng.2653.

55. Love, M.I., Huber, W., and Anders, S. (2014). Moderated estimation of fold change and dispersion for RNA-seq data with DESeq2. Genome Biol. 15, 550. 10.1186/s13059-014-0550-8.

56. Korotkevich, G., Sukhov, V., and Sergushichev, A. (2019). Fast gene set enrichment analysis. Preprint at bioRxiv, 10.1101/060012 10.1101/060012.

57. The Gene Ontology Consortium, Aleksander, S.A., Balhoff, J., Carbon, S., Cherry, J.M., Drabkin, H.J., Ebert, D., Feuermann, M., Gaudet, P., Harris, N.L., et al. (2023). The Gene Ontology knowledgebase in 2023. Genetics *224*, iyad031. 10.1093/genetics/iyad031.

58. Ashburner, M., Ball, C.A., Blake, J.A., Botstein, D., Butler, H., Cherry, J.M., Davis, A.P., Dolinski, K., Dwight, S.S., Eppig, J.T., et al. (2000). Gene Ontology: tool for the unification of biology. Nat. Genet. 25, 25–29. 10.1038/75556.

59. Kuleshov, M.V., Jones, M.R., Rouillard, A.D., Fernandez, N.F., Duan, Q., Wang, Z., Koplev, S., Jenkins, S.L., Jagodnik, K.M., Lachmann, A., et al. (2016). Enrichr: a comprehensive gene set enrichment analysis web server 2016 update. Nucleic Acids Res. 44, W90–97. 10.1093/nar/gkw377.

60. Keenan, A.B., Torre, D., Lachmann, A., Leong, A.K., Wojciechowicz, M.L., Utti, V., Jagodnik, K.M., Kropiwnicki, E., Wang, Z., and Ma’ayan, A. (2019). ChEA3: transcription factor enrichment analysis by orthogonal omics integration. Nucleic Acids Res. 47, W212–W224. 10.1093/nar/gkz446.

61. Sastry, A.V., Poudel, S., Rychel, K., Yoo, R., Lamoureux, C.R., Chauhan, S., Haiman, Z.B., Bulushi, T.A., Seif, Y., and Palsson, B.O. (2021). Mining all publicly available expression data to compute dynamic microbial transcriptional regulatory networks. Preprint at bioRxiv, 10.1101/2021.07.01.450581 10.1101/2021.07.01.450581.

62. Ester, M., Kriegel, H.-P., Sander, J., and Xu, X. (1996). A density-based algorithm for discovering clusters in large spatial databases with noise. In Proceedings of the Second International Conference on Knowledge Discovery and Data Mining KDD’96. (AAAI Press), pp. 226–231.

63. Sastry, A.V., Yuan, Y., Poudel, S., Rychel, K., Yoo, R., Lamoureux, C.R., Li, G., Burrows, J.T., Chauhan, S., Haiman, Z.B., et al. (2024). iModulonMiner and PyModulon: Software for unsupervised mining of gene expression compendia. PLOS Comput. Biol. 20, e1012546. 10.1371/journal.pcbi.1012546.

64. Pedregosa, F., Varoquaux, G., Gramfort, A., Michel, V., Thirion, B., Grisel, O., Blondel, M., Prettenhofer, P., Weiss, R., Dubourg, V., et al. (2011). Scikit-learn: Machine Learning in Python. J. Mach. Learn. Res. 12, 2825–2830.

